# Sonification of Elephant Infrasound

**DOI:** 10.64898/2026.07.07.736953

**Authors:** Arda Bozdogan, Ronald M. Aarts

**Affiliations:** Department of Electrical Engineering, Eindhoven University of Technology (TU/e), 5600 MB Eindhoven, The Netherlands

**Keywords:** psychoacoustics, digital signal processing, harmonic generation, missing fundamental, frequency estimation, infrasound, elephant communication, infrasound sonification, seismic sonification

## Abstract

Elephants and other large mammals produce low-frequency vocalizations extending well below the 20 Hz lower limit of human hearing, a regime known as infrasound. These *rumbles* serve vital social and reproductive functions over distances of several kilometers, yet they are inaudible to human observers and cannot be reproduced by conventional small loudspeakers. We present a complete signal-processing pipeline that renders sub-20 Hz elephant rumbles perceptible through a small loud-speaker by exploiting the *missing-fundamental* psychoacoustic effect. Butterworth bandpass filters isolate the infrasonic content; a full-wave integrator nonlinear device (NLD) generates the harmonic series required for virtual pitch perception; and a hysteresis-comparator fundamental-frequency estimator normalizes the NLD output. The pipeline was validated on African elephant field recordings and deployed on a credit-card-sized, low-cost single-board computer with an infrasound microphone and a small Bluetooth loudspeaker, demonstrating live operation in the field. The processed output shows a 10 dB to 15 dB elevation in the loudspeaker’s efficient band during call segments compared with background. The system enables zoo visitors and wildlife observers to perceive elephant rumbles in real time, opening new avenues for behavioral studies and public engagement with animal communication.

## 1 Introduction

Large mammals possess remarkable acoustic capabilities that extend far beyond the limits of human hearing. The elephant in particular has long been recognized as one of the most acoustically sophisticated terrestrial animals, capable of generating, detecting, and interpreting sounds across an exceptionally wide frequency range (Garstang, 2015). Yet the most ecologically significant portion of the elephant’s vocal repertoire—the low-frequency *rumble*—lies outside the audible bandwidth of the human ear and cannot be faithfully reproduced by the compact loudspeakers commonly available in field or zoo settings. This paper describes a psychoacoustic signal-processing system that bridges this perceptual gap, enabling human observers to hear elephant rumbles through a small loudspeaker in real time.

### 1.1 Infrasound in Large Mammals

Infrasound is conventionally defined as acoustic energy at frequencies below 20 Hz, the nominal lower boundary of human auditory sensitivity (Garstang, 2015). Long before the biological role of infrasound was appreciated, its use by oceanic species such as the fin whale (*Balaenoptera physalus*) had been proposed on theoretical grounds: the extreme efficiency of low-frequency propagation in water suggested that whale calls might travel thousands of kilometers (McComb et al., 2003). The discovery that terrestrial mammals could also exploit infrasound came only in the mid-1980s, when Payne et al. (1986) reported infrasonic vocalizations from captive Asian elephants (*Elephas maximus*) at the Washington Park Zoo, Portland. Most of the recorded calls ranged from 14 Hz to 24 Hz, with durations of 10 s to 15 s and sound-pressure levels of 85 dB to 90 dB (re 20 µPa) at 5 m (Payne et al., 1986). This landmark study established elephants as the first terrestrial mammals known to produce infrasound.

The vocal communication of both African savannah elephants (*Loxodonta africana*) and Asian elephants spans a rich repertoire of call types—trumpets, roars, chirps, and rumbles—but it is the rumble that carries infrasonic content and dominates long-distance signalling (Nair et al., 2009; Stoeger, 2021). Rumble fundamental frequencies typically lie between 10 Hz to 30 Hz for both species (Nair et al., 2009; Stoeger, 2021), with harmonics extending to several hundred hertz (Stoeger, 2021). Under specific physiological and environmental conditions, very large individuals may produce calls as low as 10 Hz (Garstang, 2015).

Giraffes (*Giraffa camelopardalis*) represent another large mammal whose acoustic repertoire has received attention in the context of infrasound. Although it was long assumed that giraffes, like elephants, communicated via infrasonic calls, systematic nocturnal recordings across three European zoos revealed sustained, harmonically structured *humming* vocalizations that did not reach the infrasonic domain (Baotic et al., 2015). Baotic et al. (2015) concluded that the assumption of infrasonic giraffe communication requires further investigation, highlighting the importance of rigorous acoustic measurement over anecdotal report. By contrast, the infrasonic nature of elephant rumbles is now firmly established by decades of field and laboratory work (Payne et al., 1986; Nair et al., 2009; Stoeger, 2021; Garstang, 2015; Limberger et al., 2026).

### 1.2 Acoustic Structure and Function of Elephant Rumbles

Elephant rumbles exhibit pronounced structural variation. Fundamental frequencies show frequency glides—gradual descents or ascents across the duration of a call—and harmonics are present at integer multiples of the fundamental (Stoeger, 2021; Nair et al., 2009). The social contexts in which rumbles occur span nearly the full range of elephant life: close-range affiliative interactions, herd assembly by matriarchs, contact calling within and between family groups, reproductive advertisement, and responses to disturbance or predation threat (Nair et al., 2009; Garstang, 2015).

The propagation properties of low-frequency sound make rumbles particularly suited to long-distance communication. Atmospheric attenuation is strongly frequency-dependent, scaling roughly as frequency squared for viscothermal losses (McComb et al., 2003). Infrasonic components therefore survive propagation over distances at which higher-frequency harmonics have been degraded into background noise. McComb et al. (2003) demonstrated through long-distance playback experiments that female African elephants can recognize contact calls at 2.5 km, though recognition is more reliable at 1 km to 1.5 km. Garstang (2015) used atmospheric modelling to predict that, under favorable stratification conditions, elephant calls may carry over distances exceeding 10 km.

Beyond the airborne acoustic channel, elephant vocalizations couple into the ground and propagate as seismic surface waves (O’Connell-Rodwell, 2007). O’Connell-Rodwell et al. (2006) demonstrated through field playback experiments that African elephant breeding herds respond to seismically transmitted alarm vocalizations, providing the first evidence of seismic reception in a large mammal. Elephants are hypothesized to detect ground-borne vibrations through both bone conduction via the feet and somatosensory reception, mediated by the Meissner and Pacinian corpuscles concentrated in the sole of the foot (O’Connell-Rodwell, 2007). The seismo-acoustic duality of elephant rumbles adds a further layer of complexity to their communicative repertoire and underlines the importance of multi-modal sensing approaches.

Vocal creativity and individuality are also documented in the elephant repertoire. Stoeger et al. (2021) showed that African savannah elephants develop idiosyncratic sound-production mechanisms—including nasal tissue vibration driven by ingressive airflow at the trunk tip and contraction of superficial muscles at the trunk base—that generate sounds unique to each individual. This plasticity is consistent with vocal learning abilities that parallel, in certain respects, those found in humans. Beyond individuality, the acoustic structure of the rumble also encodes information about the caller. Stoeger et al. (2014) showed that acoustic parameters extracted from low-frequency rumbles recorded under field conditions allow the age group of free-ranging African elephants to be estimated, reaching up to 70 % correct classification into four age groups and 95 % when categorising into two. This demonstrates that ecologically relevant information—such as caller age—is embedded in the very infrasonic frequency band that the present system renders audible, underlining the value of making rumbles perceptible for behavioural and conservation studies.

### 1.3 Monitoring Rumble Activity in Captive Populations

Recent work has extended infrasound research into zoo environments, where non-invasive sensing provides a means to monitor elephant behavior and welfare continuously. Limberger et al. (2026) deployed co-located seismic and infrasound sensors at the Opel-Zoo near Frankfurt, identifying over 1350 rumbles from August 2024 recordings. Call durations ranged from 1 s to 8 s with fundamentals between 10 Hz to 25 Hz, consistent with field observations. A systematic nocturnal housing schedule produced predictable increases in rumbling activity every second night, and many rumbles occurred in rapid sequences, suggesting inter-elephant interaction or external triggers. Convolutional neural networks (CNNs) trained on spectrogram images of seismic and infrasound signals achieved up to 98% classification accuracy for rumble versus noise, illustrating the feasibility of automated, continuous monitoring (Limberger et al., 2026).

### 1.4 The Perceptual and Technical Problem

Despite the ecological significance of elephant infrasound, two obstacles prevent human observers from perceiving it directly.

First, the human auditory system is insensitive below approximately 20 Hz. Simple amplification of infrasonic recordings is therefore ineffective: no matter how loud the playback, the spectral content below the audiogram threshold remains inaudible.

Second, small loudspeakers—the class of device most available in field, zoo, and portable contexts—are physically unable to radiate acoustic power efficiently at infrasonic frequencies: a compact driver’s resonance frequency is fixed by an unavoidable trade-off between cone area, moving mass, and enclosure stiffness, which keeps it stiffness-controlled, and hence acoustically inefficient, well above the elephant’s infrasonic fundamentals (Larsen and Aarts, 2002, 2004; Aarts, 2006). Infrasonic content below the loudspeaker cutoff frequency *f*_*l*_ is therefore reproduced at negligible acoustic power.

### 1.5 Psychoacoustic Approach: The Missing Fundamental

Larsen and Aarts (2004) describe several psychoacoustic methods for making low-frequency content perceptible through bandwidth-limited transducers, including frequency doubling and virtual-pitch enhancement. The present work employs the *missing-fundamental* effect, also termed *virtual pitch*. When the auditory system is presented with a harmonic series {*f*_0_, 2 *f*_0_, 3 *f*_0_, …} but the fundamental *f*_0_ itself is absent from the stimulus, the pitch perceived by the listener corresponds to *f*_0_, not to any of the physically present harmonics (Larsen and Aarts, 2004). This phenomenon arises from nonlinear processing in the peripheral auditory system and has been exploited in audio reproduction since the earliest days of broadcasting.

The key insight for the present application is that the loudspeaker need not reproduce the infrasonic fundamental directly. If a nonlinear device (NLD) generates harmonics of the elephant rumble at frequencies above the loudspeaker cutoff *f*_*l*_, the auditory system will reconstruct the percept of the original low fundamental. The temporal and harmonic structure of the call—the ecological information carried by the rumble—is thereby preserved in the perceptual domain even though no energy below *f*_*l*_ is physically emitted.

### 1.6 Contribution and Outline

This paper makes four main contributions. First, we present a complete mono-channel psychoacoustic pipeline—filter design, NLD, and a fundamental-frequency estimator with a time-varying extension—optimized for sub-20 Hz elephant rumbles and a 90 Hz-cutoff small loudspeaker. Second, we validate the pipeline on African elephant field recordings, demonstrating a 10 dB to 15 dB elevation in the loud-speaker’s efficient band during call segments. Third, we provide an embedded real-time implementation on a credit-card-sized, low-cost single-board computer, offering live sonification of elephant infrasound on portable, field-deployable hardware. Finally, the resulting system enables wildlife and zoo observers to perceive elephant infrasound in real time, supporting behavioral studies and public engagement with animal communication.

The remainder of the paper is organized as follows. Section 2 covers the theoretical background on infrasound and psychoacoustics. Section 3 describes the signal-processing pipeline. Section 4 outlines the evaluation methodology. Section 5 presents results on field recordings. Section 6 describes the embedded deployment. Section 7 discusses findings and limitations, and Section 8 concludes.

## 2 Theoretical Background

### 2.1 Infrasound and Human Auditory Limits

The International Organization for Standardization defines the audible frequency range as 20 Hz to 20000 Hz, placing infrasound below this lower bound. Human audiometric sensitivity declines steeply below 50 Hz; at 20 Hz the threshold of hearing is approximately 74 dB SPL, compared with 0 dB SPL at the frequency of maximum sensitivity around 3 kHz. Below 20 Hz physical stimulation of the cochlea is possible only at very high sound levels, and below about 10 Hz the sensation becomes predominantly tactile rather than auditory.

Elephants are far better equipped to perceive infrasound: the female Asian elephant has been shown to detect a 60 dB signal at frequencies as low as 17 Hz and as high as 10.5 kHz, the widest known hearing range of any non-human mammal (Garstang, 2015).

### 2.2 Missing Fundamental and Virtual Pitch

The missing-fundamental effect is described in detail by Larsen and Aarts (2004). When a listener is presented with harmonics {2 *f*_0_, 3 *f*_0_, 4 *f*_0_, …} of a fundamental *f*_0_ that is physically absent from the stimulus, the auditory system infers a pitch corresponding to *f*_0_, as illustrated schematically in Fig. 1. The mechanism is attributed to nonlinear distortion in the inner ear and pattern-matching processes in the auditory cortex. For the purpose of small loudspeaker enhancement, the missing fundamental eliminates the requirement for the transducer to reproduce the lowest frequency component: as long as a sufficient number of harmonics above *f*_*l*_ are present, the listener will perceive the correct pitch. That the auditory system extracts the fundamental of a complex harmonic sound accurately, and not merely approximately, is well established psychoacoustically. Le Goff et al. (2004) measured thresholds for detecting mistuning of the fundamental component in harmonic complex tones with fundamentals as low as 60 Hz, showing that listeners are sensitive to small deviations in the low-frequency fundamental even when it is embedded in, or reconstructed from, the surrounding harmonics. This quantitative evidence underpins the reliability of the virtual-pitch percept exploited here and removes the need to treat perception of the synthesized rumble pitch as merely anecdotal.

**Figure 1:**
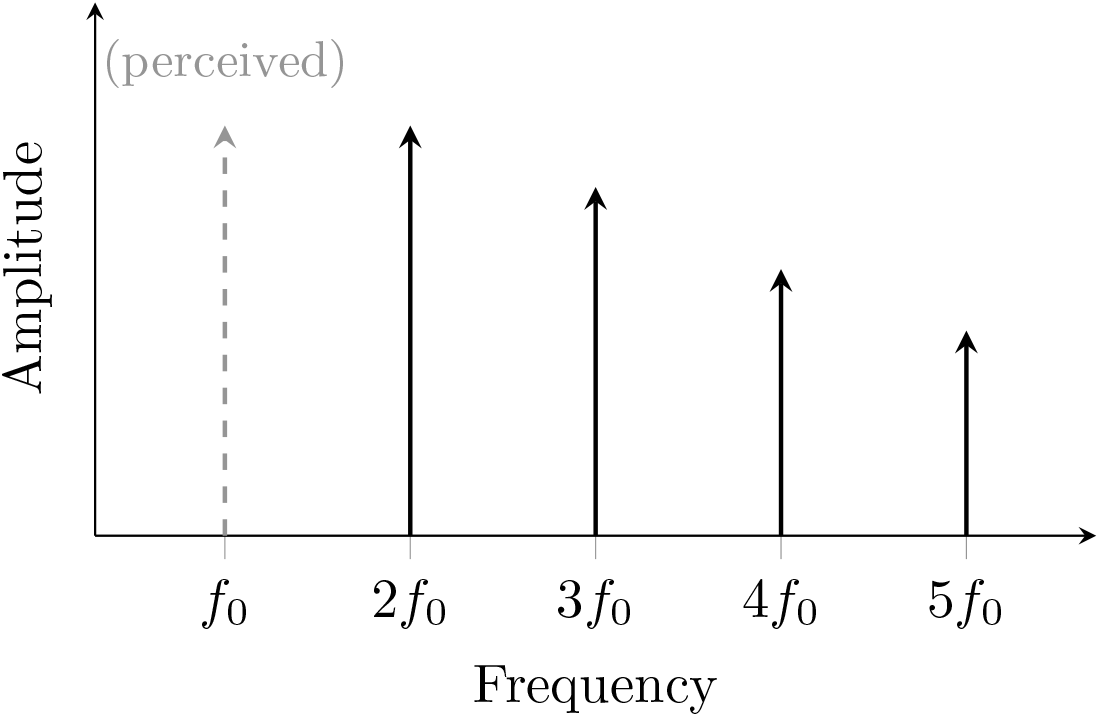
Illustration of the missing fundamental effect. The dashed component at *f*_0_ is not physically reproduced; its pitch is perceived by the auditory system from the harmonics at 2 *f*_0_, 3 *f*_0_, 4 *f*_0_, …

### 2.3 Loudspeaker Physics and the Cutoff Frequency

As outlined in Section 1 and detailed in Larsen and Aarts (2004), a moving-coil loudspeaker has a resonance frequency *f*_*l*_ = *ω*_0_/(2*π*) below which radiated acoustic power drops precipitously. In the stiffness-controlled region (*f < f*_*l*_) cone excursion is large but poorly coupled to the surrounding medium; in the mass-controlled region (*f > f*_*l*_) excursion falls as 1*/ω*^2^ but acoustic radiation efficiency rises. One octave above *f*_*l*_ the driver already operates substantially more efficiently and linearly. The psychoacoustic approach in this paper exploits this regime by placing all synthesized harmonics above *f*_*l*_.

## 3 Signal Processing Pipeline

The pipeline, illustrated schematically in Fig. 2, is a mono-channel digital processing scheme that extends the stereo architecture described by Larsen and Aarts (2004) to the specific requirements of elephant infrasound sonification.

**Figure 2:**
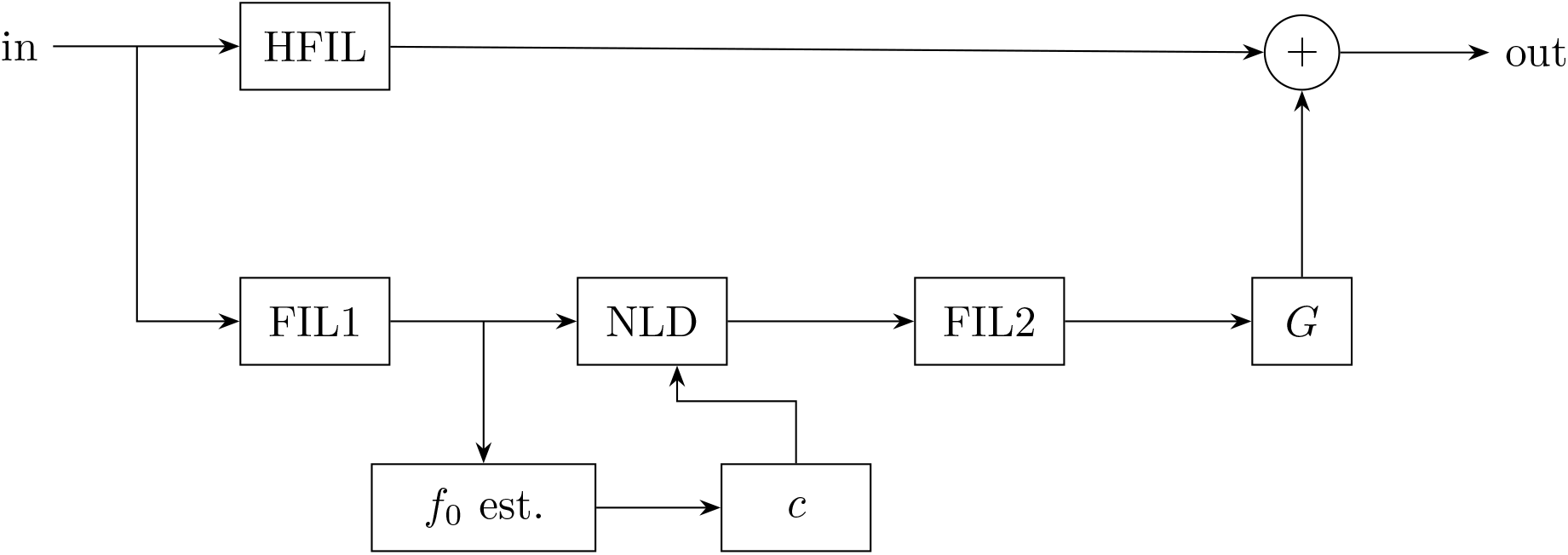
Signal processing scheme, adapted from Larsen and Aarts (2004) and modified for sonification of elephant infrasound.

### 3.1 Filter Design

Three fourth-order Butterworth filters are used. Butterworth filters are chosen for their maximally flat passband response (Larsen and Aarts, 2004), which avoids coloring the harmonic content presented to the NLD.

The cutoff frequencies are dictated by the loudspeaker physics described in Section 2. For the small loudspeaker used in this work, the manufacturer reports *f*_*l*_ ≈ 91 Hz, rounded to *f*_*l*_ = 90 Hz for design purposes. The three filters are:

- **hfil**: High-pass at 90 Hz. Preserves audible harmonic content already present in the elephant signal and passes it directly to the output.
- **fil****1**: Bandpass from 10 Hz to 90 Hz. Isolates the infrasonic fundamental content. The 10 Hz lower bound covers the full reported range of elephant rumble fundamentals (Nair et al., 2009; Stoeger, 2021). The passband of approximately 3.2 octaves marginally exceeds the three-octave guideline of Larsen and Aarts (2004) above which intermodulation distortion at the NLD output becomes perceptible; listening confirmed that no audible artifacts arose for the recordings used in this work.
- **fil****2**: Bandpass from 90 Hz to 180 Hz, corresponding to [*f*_*l*_, 2 *f*_*l*_]. The lower bound excludes harmonics below the efficient radiation band; the upper bound restricts the harmonic set to a single octave, avoiding the sharp timbre that a very broad harmonic spectrum produces (Larsen and Aarts, 2004).

### 3.2 Fundamental Frequency Estimation

Accurate knowledge of *f*_0_ is required solely to set the normalization constant *c* of the NLD (Section 3.3); the NLD reset timing is governed by zero crossings and is independent of the estimate. The estimator is a software hysteresis comparator operating on a 10 Hz to 30 Hz bandpass pre-filtered version of the fil1 output, which isolates the fundamental band and suppresses higher harmonics that would otherwise corrupt the period measurement.

Because elephant rumbles exhibit a frequency glide rather than a stationary fundamental, a time-varying extension is implemented: *f*_0_ is estimated over successive 2 s windows using an RMS-based activity detector that suppresses estimation during background noise segments. The normalization constant *c* is recomputed at each window boundary and linearly interpolated to sample resolution between consecutive windows, ensuring smooth gain variation without discontinuities.

Alternatively, the fundamental could be tracked using a highly efficient frequency tracker, as described by Aarts (2021).

### 3.3 Nonlinear Device

The NLD generates the harmonic series required for the missing-fundamental effect. Following Larsen and Aarts (2004), a *full-wave integrator*—which accumulates the absolute value of the input signal and resets to zero at each positive-going zero crossing—is used in preference to a full-wave rectifier, since it produces all odd and even harmonics of *f*_0_ (with amplitudes decaying as 1*/n*^2^ for harmonic index *n*) rather than only the even harmonics obtained from a rectifier, yielding more robust pitch perception; the full mathematical treatment is given by Larsen and Aarts (2004). The constant *c* scales the NLD output to maintain perceptual consistency across recordings at different amplitude levels.

The gain stage *G* at the output compensates for the 1*/n*^2^ harmonic rolloff: because the elephant fundamental is well below *f*_*l*_, fil2 selects relatively high-order harmonics whose amplitudes are accordingly small. Because this NLD is linear in its amplitude behavior, it preserves the original envelope shape of the time signal.

## 4 Evaluation Methodology

Evaluation used African elephant field recordings provided by Angela Stöger (University of Vienna) and Fabian Limberger (Goethe University Frankfurt), together with recordings made at the Opel-Zoo Kronberg (Limberger et al., 2026) and at the Eindhoven Zoo (Beekse Bergen) using a Jordan ATD4-S infrasound microphone (Dr. Jordan Design, 2024). The recordings span sampling rates of 8000 Hz and higher, with infrasonic content verified by visual inspection of spectrograms in the 0 Hz to 200 Hz range (Hamming window, 8192 samples, 7168-sample overlap, 16384-point FFT).

Performance is assessed by:

1. **Spectral elevation**: The power spectral density (PSD) during a call segment is compared with a background reference; a consistent elevation in the fil2 passband (90 Hz to 180 Hz) confirms that the pipeline delivers energy into the loudspeaker’s efficient band in response to the call.
2. **Listening test**: A human listener verifies whether audible low-frequency content is present during call segments in the processed output and absent in the unprocessed signal, a judgement supported by established psychoacoustic thresholds for perceiving the fundamental of low-frequency harmonic complexes (Le Goff et al., 2004).
3. **Filter configuration comparison**: The 90 Hz configuration (*f*_*l*_ = *f*_*c*_ = 90 Hz) is compared with a 40 Hz alternative to confirm the design rationale.

## 5 Results

A representative result is presented for recording 02T19 from the Limberger dataset. The fundamental frequency was estimated at *f*_0_ = 18.10 Hz by the hysteresis comparator. A descending frequency trace near *f*_0_, with a second harmonic at approximately 2 *f*_0_, confirms the presence of the call, rising sharply in intensity during the call segment (approximately 8 s to 15 s; Fig. 3).

**Figure 3:**
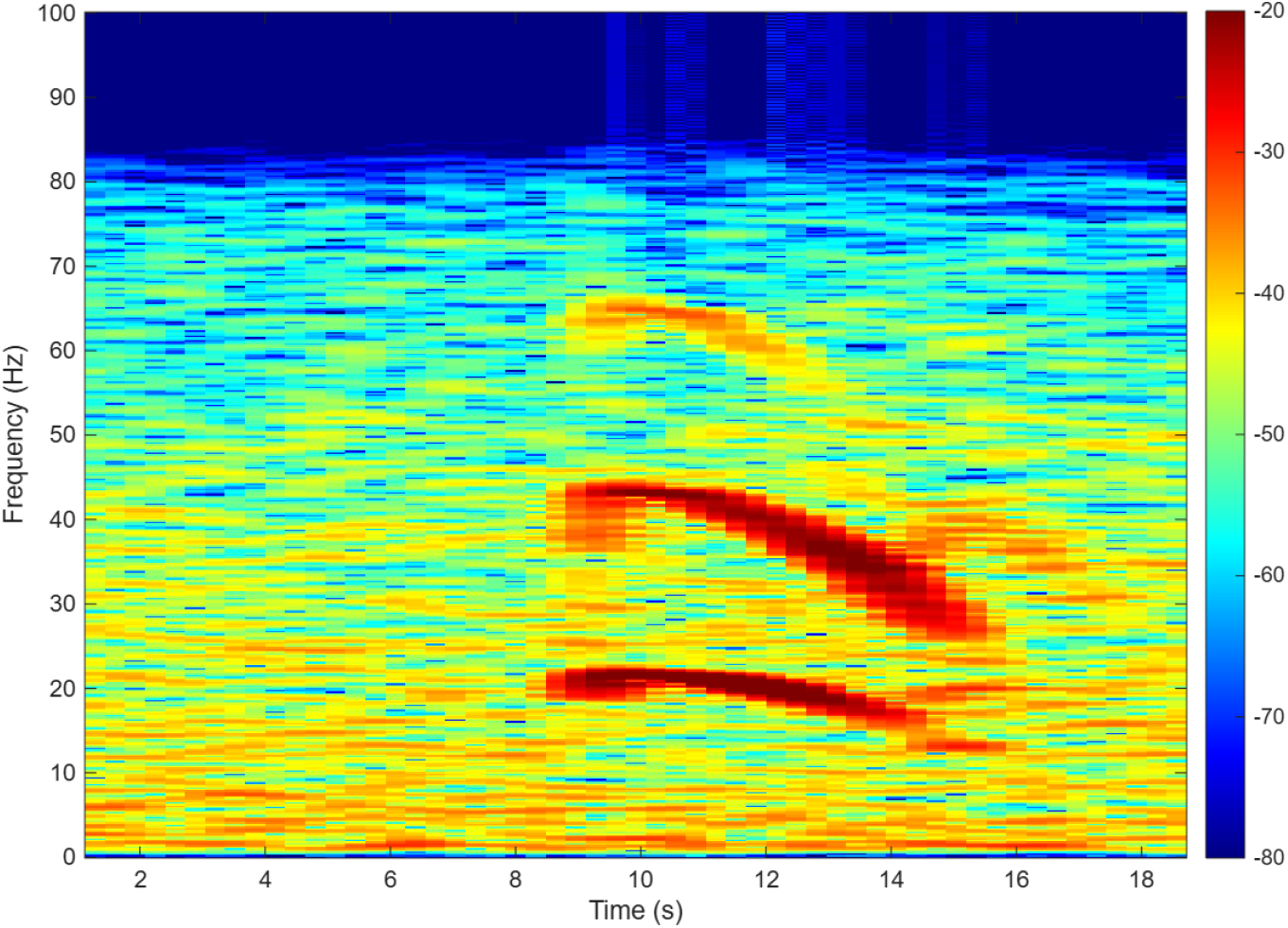
Raw spectrogram of recording 02T19. A descending fundamental trace near *f*_0_ = 18.10 Hz and its second harmonic are visible during the call segment (approximately 8 s to 15 s).

During the call segment, a stack of horizontal harmonic bands extends from approximately 32 Hz to 180 Hz, with bands above 90 Hz corresponding to synthesized harmonics within the fil2 passband; before and after the call, this harmonic structure collapses, confirming that the output is driven by the elephant call rather than background noise (Fig. 4).

**Figure 4:**
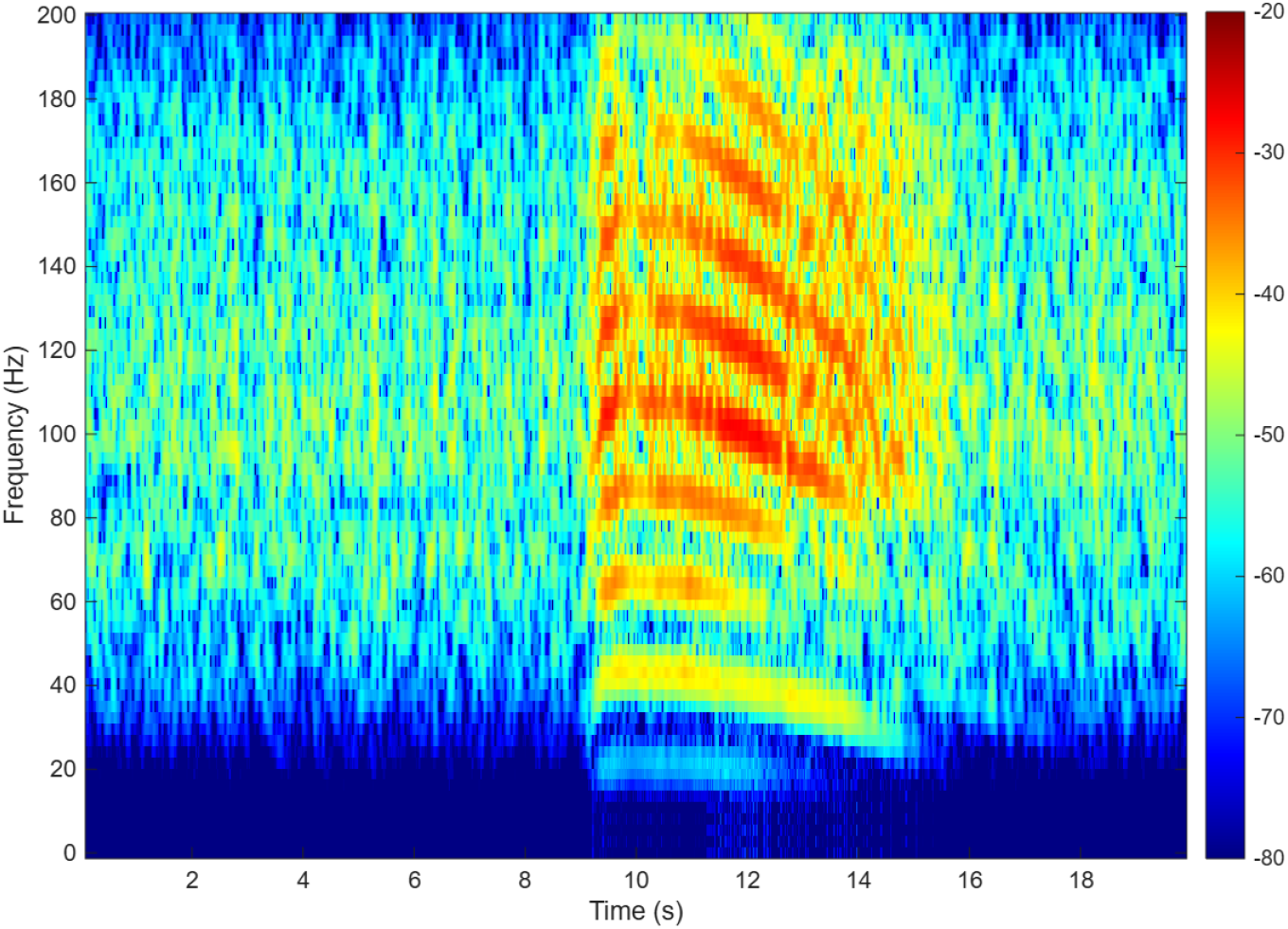
Processed (sonified) spectrogram of recording 02T19. Synthesized harmonics fill the 32 Hz to 180 Hz band during the call segment and collapse outside it, confirming that the audible output tracks the elephant call.

To assess consistency across calls, the same processing was applied to two further recordings from the Limberger dataset, 19T01 and 26T19, both of which exhibit the characteristic paired fundamental-plus-second-harmonic glide of an elephant rumble (Fig. 5). In both cases, the processed output reproduces a corresponding stack of harmonic bands above 90 Hz that is present only during the call segment and absent from the surrounding background (Fig. 6). The harmonic structure is somewhat weaker and more fragmented for 26T19 than for 02T19 or 19T01, consistent with its lower raw-spectrogram call intensity, but the call-aligned elevation above *f*_*l*_ is clearly visible in all three recordings.

**Figure 5:**
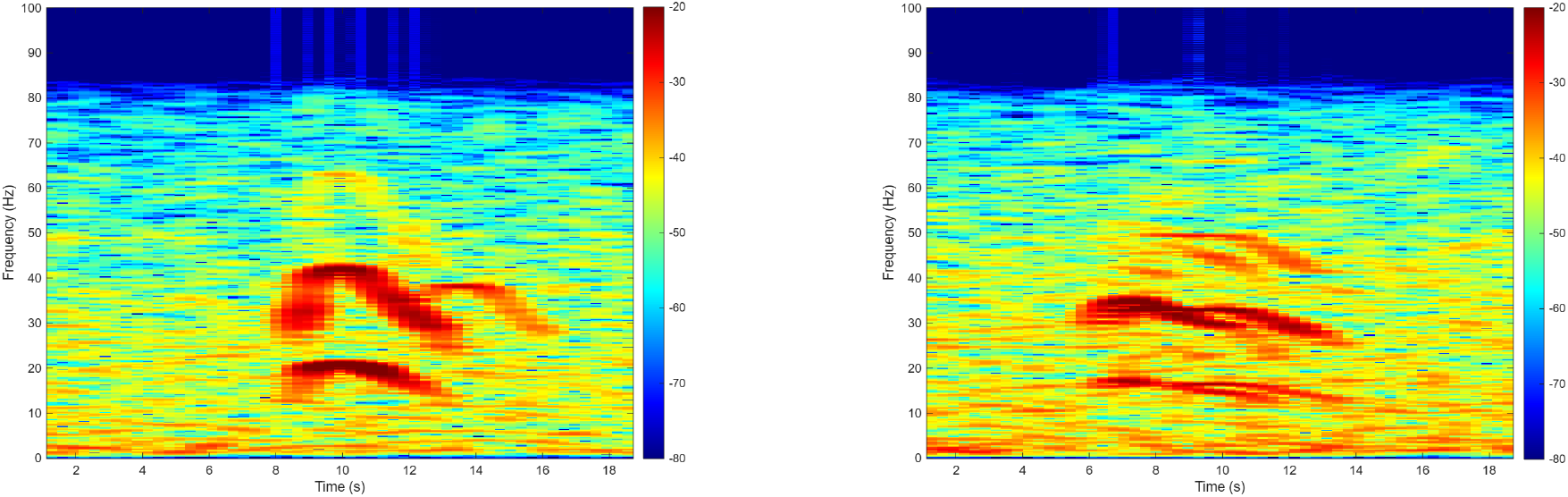
Raw spectrograms of two further recordings from the Limberger dataset, before processing: 19T01 (left), with a call segment around 8 s to 13 s, and 26T19 (right), with a call segment around 6 s to 13 s.

**Figure 6:**
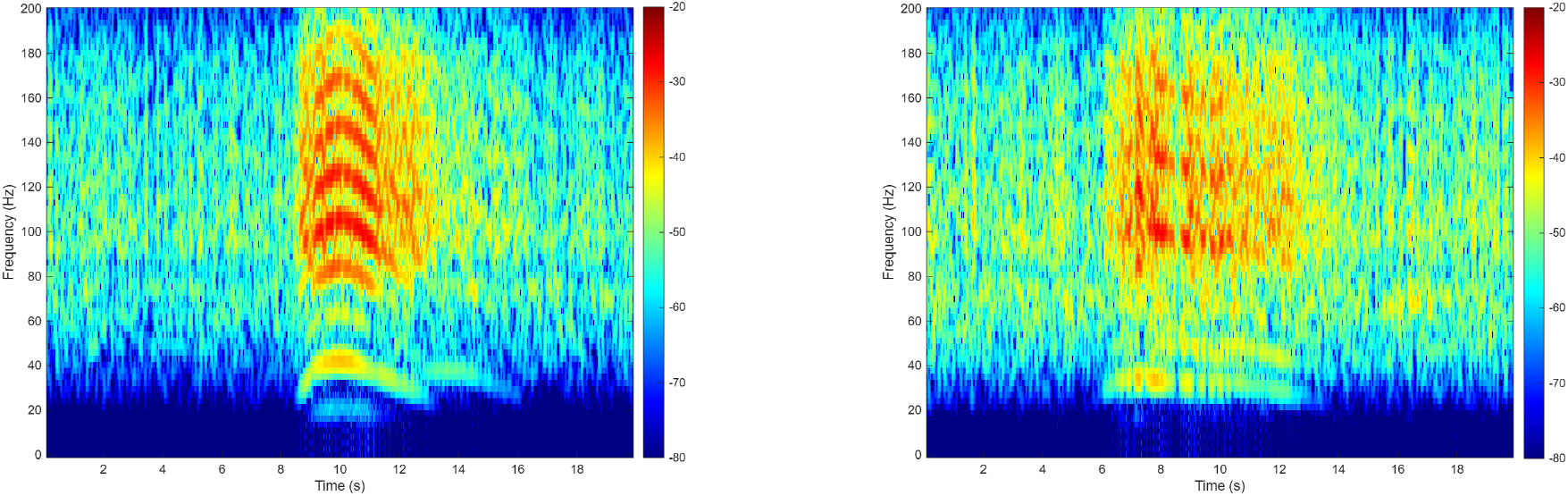
Spectrograms of the same two recordings after sonification: 19T01 (left) and 26T19 (right). Synthesized harmonic energy is concentrated above 90 Hz during the call segment only, replicating the pattern observed for recording 02T19 (Figs. 3–4).

The PSD comparison in the fil2 passband shows a 10 dB to 15 dB elevation during the call segment relative to the background. The processing path contributes demonstrably above 90 Hz: the full processed output exceeds the hfil direct path alone by this margin.

For the filter configuration comparison, setting *f*_*c*_ = 40 Hz concentrates synthesized energy in the 40 Hz to 90 Hz band where the small loudspeaker used in this work radiates poorly, yielding a 10 dB to 15 dB deficit above 90 Hz relative to the 90 Hz configuration. This confirms that matching *f*_*c*_ to the measured loudspeaker cutoff *f*_*l*_ is essential for efficient sonification.

Listening confirmed clearly audible low-frequency content during call segments that was absent in the unprocessed signal. The temporal structure of the rumble—onset, glide, and decay—was readily perceptible in the processed output. This outcome is consistent with the published psychoacoustic evidence that the auditory system reliably recovers the fundamental of a low-frequency harmonic complex (Le Goff et al., 2004). Furthermore, because the signal envelope is preserved by the NLD, the envelope of the rumble remains close to its original shape, but is now audible.

## 6 Embedded Deployment

The MATLAB offline pipeline was ported to Python and deployed on a Raspberry Pi 4B—a low-cost, credit-card-sized single-board computer developed by the Raspberry Pi Foundation, carrying a quad-core processor, up to 8 GB of RAM, and USB, Bluetooth, and Wi-Fi connectivity on a single board—to demonstrate live, portable, field-deployable operation on embedded hardware. The hardware chain consists of the infrasound microphone connected via USB and a small loudspeaker connected via Bluetooth.

The embedded system continuously listens to short segments of audio, passes each segment through the psychoacoustic pipeline described above, and plays the result back with minimal delay, so that a sonified rumble is heard shortly after it occurs. A simple safeguard keeps the output level from distorting when the processed signal would otherwise be too loud for the loudspeaker.

The system was verified using a synthetic 15 Hz sine wave, confirming audible output through the small loudspeaker. The implementation is publicly available at https://github.com/ArdaBozdogan/Elephant-Infrasound-BEP.

## 7 Discussion

The pipeline successfully produces perceptible low-frequency output from elephant rumble recordings through a small loudspeaker, validating the core hypothesis: the missing-fundamental effect, as described by Larsen and Aarts (2004), is applicable to sub-20 Hz animal vocalizations.

Virtual-pitch perception of low-frequency harmonic complexes is itself well established: listeners reliably extract, and are finely sensitive to, the fundamental of such complexes down to at least 60 Hz (Le Goff et al., 2004; Larsen and Aarts, 2004). The present application extends this principle to the sub-20 Hz fundamentals of elephant rumbles, using relatively high-order harmonics given the low fundamental.

The primary technical limitation is the breakdown of the NLD’s near-sinusoidal input assumption under broadband field noise. A real rumble distributes energy across the full fil1 passband rather than concentrating it near *f*_0_, causing the integrator to produce a non-periodic output. The 10 Hz to 30 Hz prefilter for the estimator is a practical mitigation but not a general solution: for fundamentals below 15 Hz the second harmonic re-enters the pre-filter passband. More robust frequency estimation methods—autocorrelation, cepstral analysis, or the recursive sample-by-sample tracker described by Larsen and Aarts (2004)—represent natural directions for future work.

From the perspective of behavioral science and conservation, the system opens several practical applications. In real-time field and zoo observation, wildlife rangers and zoo visitors can perceive elephant rumbles live, without specialized training or instrumentation, while simultaneously observing the animal’s behavior. For behavioral-correlate research, the real-time coupling of auditory percept to visible behavior enriches the observational experience and may facilitate the detection of behavioral correlates of specific call types. Combined with automated classification systems of the kind developed by Limberger et al. (2026), the pipeline could also support captive welfare monitoring, forming part of a broader welfare-monitoring platform for captive elephant populations. More broadly, rendering an otherwise inaccessible communication channel audible supports public engagement and education, allowing non-specialist audiences to engage directly with a sensory modality that elephants use but humans cannot otherwise perceive. Finally, the same missing-fundamental pipeline is not restricted to airborne micro-phone signals: elephant rumbles also couple into the ground as seismic surface waves, and locomotion or trampling produces additional motion-induced ground vibrations that are not captured by infrasound sensors at all (Limberger et al., 2026; O’Connell-Rodwell, 2007). Co-located geophone recordings, such as those collected alongside infrasound microphones by Limberger et al. (2026), are themselves sub-20 Hz time series and can be passed through the identical filter–NLD–estimator chain to render these ground vibrations audible, extending the system from a purely acoustic tool toward a multi-modal (acoustic-and-seismic) sonification platform.

## 8 Conclusion

We have presented a psychoacoustic signal-processing system that renders sub-20 Hz elephant rumbles audible through a small loudspeaker by exploiting the missing-fundamental effect (Larsen and Aarts, 2004). The pipeline comprises Butterworth bandpass filters, a full-wave integrator nonlinear device, and a time-varying hysteresis-comparator fundamental-frequency estimator. Validation on African elephant field recordings demonstrates a 10 dB to 15 dB elevation in the loudspeaker’s efficient radiation band during call segments. An embedded deployment on a credit-card-sized, low-cost single-board computer demonstrates that the pipeline can run as a live sonification system on portable, field-deployable hardware.

The system addresses a practical perceptual gap: elephant infrasound is ecologically and socially significant (Payne et al., 1986; McComb et al., 2003; O’Connell-Rodwell et al., 2006; O’Connell-Rodwell, 2007; Garstang, 2015; Nair et al., 2009; Stoeger, 2021; Limberger et al., 2026), yet invisible to the unaided human ear and inaudible through conventional small loudspeakers. By making these calls perceptible in real time, the system enables wildlife observers and zoo visitors to experience elephant communication directly, supporting both public engagement and scientific study of animal behavior.

Future work should target more robust frequency estimation under broadband field noise, a formal perceptual study to validate virtual pitch perception at the relevant harmonic indices, further reduction of system latency toward a sample-by-sample regime (Larsen and Aarts, 2004), and extension of the pipeline to co-located geophone channels so that ground-borne vibrations induced by elephant locomotion and seismic rumble components can be rendered audible alongside the airborne signal (Limberger et al., 2026).

## Acknowledgment

The authors thank Angela Stöger (University of Vienna) for providing African elephant field recordings, Fabian Limberger (Goethe University Frankfurt) for the primary test recordings; and Okke Ouweltjes (Philips Research), Aki Härmä (University of Maastricht) and Stijn Berger (Research & Animal Welfare Coordinator of Libéma) for recordings at the Eindhoven Zoo and the Beekse Bergen.

## Author Contributions

**CRediT author statement:** Ronald M. Aarts: Conceptualization, Writing – original draft, Writing – review & editing. Arda Bozdogan: Writing – original draft, Software.

## Disclosure Statement

No potential conflict of interest was reported by the author(s).

## Notes

### Competing Interest Statement

The authors have declared no competing interest.

